# Acute Increase of Excitatory Activity in Pyramidal Neurons of Rat Motor Cortex under Static Magnetic Field

**DOI:** 10.64898/2026.07.20.739680

**Authors:** Rebecca C. Klein, Stefan M. Goetz, Wolfgang B. Liedtke, Scott D. Moore, Angel V. Peterchev

**Affiliations:** Department of Psychiatry & Behavioral Sciences, Duke University, Durham, NC, USA; Durham VA Medical Center, Durham, NC, USA; Department of Neurology, Duke University, Durham, NC, USA; Department of Biomedical Engineering, Duke University, Durham, NC, USA; Department of Electrical & Computer Engineering, Duke University, Durham, NC, USA; Department of Neurosurgery, Duke University, Durham, NC, USA

**Keywords:** static magnetic field stimulation, pyramidal neuron, patch-clamp recording, excitatory post-synaptic currents, miniature excitatory post-synaptic currents, inhibitory post-synaptic currents

## Abstract

**Objectives:** Transcranial application of a static magnetic field (SMF) was reported to result in subsequent modulation of neural excitability in the human motor cortex, but the acute mechanisms underlying this effect are unknown. We explore the mechanisms of this phenomenon with patch-clamp recording in rat brain slices during SMF exposure.

**Materials and Methods:** Patch-clamp recording from layer II/III pyramidal neurons in motor cortex of acutely prepared brain slices were conducted during exposure to 0.20–0.35 T SMF or sham.

**Results:** During SMF exposure we observed an increase in the frequency of spontaneous excitatory postsynaptic currents (sEPSCs) as well as miniature excitatory post-synaptic currents (mEPSCs) recorded in the presence of TTX to suppress action potentials and thus also network effects. There was a significant acute increase in sEPSC frequency for SMF exposure duration of both 6 min and 10 min, but not for 10 min sham exposure. After SMF exposure, the sEPSC and mEPSC frequency returned to baseline. The frequency of spontaneous inhibitory postsynaptic currents (sIPSCs) was unaffected by SMF. The amplitude of the postsynaptic currents decreased with time for all recordings regardless of the condition, presumably due to the expected gradual deterioration of the patch clamp seal.

**Conclusions:** The acute effect of SMF exposure on sEPSC and mEPSC frequency, but not amplitude, is consistent with the assumption of a presynaptic or synaptic site mediating the effect. Furthermore, the consistency of the effect between sEPSCs and mEPSCs suggests that the effect is not related to action potential propagation in the presynaptic axon. The effect of SMF on EPSCs and not IPSCs may be related to the larger length of excitatory axons compared to inhibitory axons, or to effects on extracellular ionic gradients within the slice.

## Introduction

Strong static magnetic field (SMF) of several hundred millitesla applied transcranially to the human primary motor cortex has been reported to reduce excitability of the corticospinal tract and alter both intracortical facilitation and inhibition (1–8). Other in-vivo observations of SMF effects include modulation of somatosensory potentials in humans (9–11), pain alleviation in mice and humans (12–15), impairment of visual processing in primates and cats (16), changes in evoked potentials and auditory processing in humans (9, 17), as well as learning and behavioral changes in rats (18).

These observations are important for at least two reasons. First, compared to other neuromodulation methods, a substantial advantage of transcranial SMF stimulation is that the requisite static field can be generated inexpensively with compact rare-earth permanent magnets, without the need for an accompanying energy source. Thus, if robust neuromodulatory effects can be achieved, such magnets can be an attractive tool for both research and therapy (19). Second, the presence of neuromodulatory SMF effects may have to be considered in functional magnetic resonance imaging (MRI) studies, where the field strength is an order of magnitude higher (typically 1.5 or 3 tesla).

However, divergent experimental observations and sensitivity to the probing parameters indicate that the effects of SMF on the brain are not yet well established or understood (4, 8, 20–25). In addition, few studies have explored the mechanisms underlying the effects of SMF on neural tissue, and a fundamental mechanism is not established. Proposed theoretical explanations include several phenomena, such as Lorentz force effects on moving charges resulting in, for example, anisotropy of ion diffusion, which is relevant for many biochemical reactions; magnetic forces on the diamagnetic lipid layers or lipid rafts of the cell membrane, possibly coupled with TRP-channel-mediated mechanoreception pathways; and influence of the magnetic flux on the potential energy profile of electrochemical reactions inside the cell due to interaction with electron spins in spin-sensitive biochemical processes (8, 26–38).

In experimental models, SMF was reported to affect the sensory nerve threshold (39, 40), conduction in the guinea pig and human spinal cord (41, 42), spiking in the lateral geniculate nucleus (43) as well as of single neurons (44, 45), the dynamics of certain ion channels (29, 46, 47), and gene expression as well as functional maturation of cultured rat hippocampal neurons (48). Furthermore, the intracellular calcium level may be sensitive to SMF as has been studied indirectly for neurons (49, 50) and imaged in astrocytes as well as lymphocytes (51, 52). Additional studies documented increased oxidative stress as well as reduced mitochondrial function in astrocytes (53) and increased calcium concentrations and calcium channel subunit expression in oligodendrocyte precursor cells (54). Finally, transient increase of Cl^−^ ion conductivity, reduction of neural excitability, and swelling of pyramidal neurons have been reported after SMF exposure of mouse brain slices (55). Still, the acute effects of SMF on the cerebral cortex are not well characterized.

To explore the acute effects of SMF exposure on neuronal activity, we performed whole-cell patch-clamp recordings of pyramidal neurons in the motor cortex of rat brain slices. We measured SMF effects of both excitatory and inhibitory transmission. Combined with the observations from prior studies, the presented data indicate that SMF could significantly influence neural activity and thus may provide practical utility in clinical applications.

## Materials and Methods

### Brain Slice Preparation

Animal procedures followed the National Research Council’s Guide for the Care and Use of Laboratory Animals. We prepared acute coronal slices with a thickness of 350 µm from 3–5 week old male Sprague Dawley rats. Rats were anaesthetized with isoflurane, decapitated, and brains were quickly removed to be placed in ice-cold oxygenated artificial cerebrospinal fluid (126 mM NaCl, 2.5 mM KCl, 1.3 mM MgCl_2_, 2.5 mM CaCl_2_, 1 mM NaH_2_PO_4_, 25 mM NaHCO_3_, and 10 mM glucose). We generated slices with a Vibratome 1000 Plus (Vibratome, Campden, England); we glued the brains to the vibratome stage and kept the brains immersed in the cooled oxygenated artificial cerebrospinal fluid. We incubated the cut slices at room temperature for at least one hour prior to electrophysiological recordings.

### Patch-clamp Recording

We located the premotor cortex (M2, Bregma 0.48 to −1.80) and performed whole-cell patch-clamp recording in layer II/III cortical pyramidal neurons. We identified the pyramidal neurons based on their morphology and firing properties. We stained the neuron recorded and confirmed morphological properties by adding 0.3% biocytin (Sigma) to the internal solution on the day of the experiment.

Spontaneous excitatory postsynaptic currents (sEPSCs), associated AMPA/Kainate responses, were isolated using patch pipettes with resistances of 3–5 MΩ filled with an internal solution containing 140 mM potassium gluconate, 0.5 mM CaCl_2_, 2 mM MgATP, 2 mM MgCl_2_, 5 mM EGTA, and 10 mM Hepes (pH 7.3). Miniature EPSCs (mEPSCs) were recorded in the presence of 1 µM tetrodotoxin (TTX), which suppresses action potentials to differentiate the role of presynaptic action potentials from other synaptic effects and network activity. We further isolated spontaneous inhibitory postsynaptic currents (sIPSCs) associated with GABA responses with an internal solution containing 140 mM CsCl, 2 mM MgATP, 2 mM MgCl_2_, 0.5 mM EGTA, and 10 mM HEPES, 4 mM Mg-ATP, 0.5 mM Tris GTP, 5 mM QX-314 (pH 7.3).

During the experiments, we perfused the slices at room temperature (22–25 °C) with oxygenated artificial cerebrospinal fluid. Neurons were clamped at –70 mV and currents were recorded using an Axopatch 200B amplifier, filtered at 1 kHz, and sampled at 10 kHz using pClamp10.1 software (Molecular Devices, Sunnyvale, CA, USA). Each trial included a measurement of the membrane properties, including resting potential, access resistance, and response to electrical depolarization (baseline, during SMF exposure, and after SMF exposure). The resting membrane potential could not be measured for sIPSCs due to the intracellular solution (CsCl) used for recording. The resting potential and access resistance did not change by more than 10% throughout the experiments for any of the recorded neurons. We discarded all neurons whose access resistance increased to more than 20 MΩ. Each measurement for SMF or Sham was a single neuron from a fresh slice. To avoid influence of light on the neural activity of the brain slices or on the metal–electrolyte interface in the pipette (56–60), which might be in sync with exposure to SMF or Sham due to their casted shadows, we kept the ambient light below 100 lx and excluded data above that limit (see Supplementary Material).

### Static Magnetic Field Exposure

A cylindrical 2”×1” NdFeB permanent magnet (grade N52, 235 lbs pull force, P/N APPLIED-MAGNETS-17, Applied Magnets, Plano, TX, USA) served for SMF exposure. A micromanipulator positioned the magnet 3.5–4 mm above the brain slice (see Fig. 1A). The south pole faced the sample. To initiate SMF stimulation the magnet was rotated approximately 90 degrees horizontally with the micromanipulator until it was positioned directly over the slice and then slowly lowered vertically. The magnet was moved slowly (∼ 3–5 °/s) to prevent electric field induction due to rapid magnetic flux changes and mechanical artifacts. The brain slice was located near the edge of the magnet cylinder to allow access of the patch pipette. An air table (Vibraplane, Kinetic Systems, Inc., Boston, MA, USA) suspended the entire setup including the slice chamber and the micromanipulator holding the magnet to reduce vibration artifacts from the magnet arm movement and from the environment. The slices were consistently oriented relative to the magnet as illustrated in Fig. 1B to control directional sources of variability. The apical dendrite of the patched pyramidal cells projected toward the surface to layer I (Fig. 1C); thus, the apical dendrites were projecting from the magnet edge towards its center.

**Figure 1.**
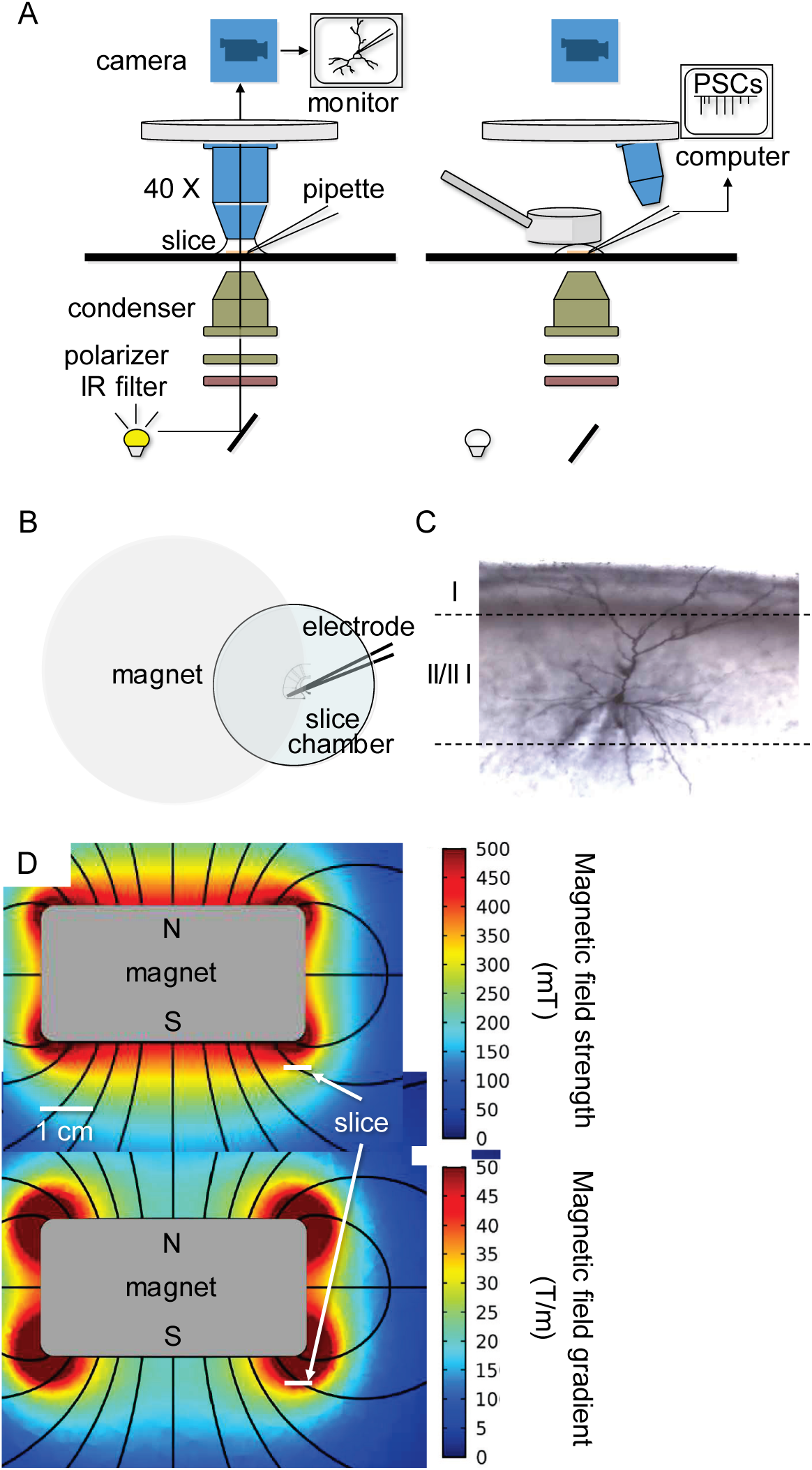
A: Schematic diagram of patch-clamp recording setup during neuron patch clamping (left) and during subsequent static magnetic field (SMF) or sham exposure and postsynaptic current (PSC) measurement (right). B: Illustration of magnet position relative to brain slice, electrode, and recording chamber. C: Representative stained layer II/III pyramidal neuron from which patch-clamp recordings were obtained. D: Position of brain slice relative to simulated distribution of magnet field strength and spatial gradient (adapted with permission from (61)).

The magnet generated a flux density of 0.20–0.35 T in the brain slice as measured with a Tesla-meter (Model 5080, F. W. Bell, Milwaukie, OR, USA). As illustrated in Fig. 1D, the magnetic field under the magnet edge was angled with respect to the slice surface. The magnetic field spatial gradient in the slice was estimated to range 35–50 T/m (Fig. 1D). A nonmagnetic aluminum cylinder identical in size and similar in color to the permanent magnet served as sham in order to reproduce various potential artifacts (e.g., mechanical, thermal, or optical reflectance) resulting from the introduction of the real magnet.

After recording of a stable baseline for at least 6 min, we exposed the slice to either SMF or sham SMF. The slice was exposed to SMF or sham for 10 min followed by continued recording after removal of the field for an additional 10 min. Access resistance and resting membrane potential were periodically measured throughout the recording between the two-minute recording bins to ensure that these values did not change ±10% over baseline measures.

### Data Analysis

The post-synaptic current spike intervals and amplitudes were measured using Mini Analysis software (Synaptosoft, Inc. Decatur, GA, USA). Average spike frequency and amplitude were computed for bins that spanned 2 min of recording before the reported time point. The average baseline was taken from the first 6 minutes of recording prior to magnet introduction and was then normalized to 100%. Averaged data are presented as baseline-normalized mean ± s.e.m.

Mixed-effect models in JMP Pro 14 (SAS Institute, Cary, NC, USA) served for the main statistical analyses with additional processing and plotting in MATLAB (The MathWorks, Natick, MA, USA). In the analyses, time point and exposure condition (SMF or sham) were treated as fixed effects and cell as a random effect nested in the exposure condition. Significant results of the mixed-effects model were followed up with post-hoc tests as follows. Time points during and following active or sham SMF exposure were compared to baseline using Dunnett’s or Dunnett–Hsu post-hoc tests. sEPSCs for SMF and sham exposure were compared for each time point with pair-wise Student’s t-tests followed by the Benjamini–Hochberg false discovery rate correction for multiple comparisons. Multiple regression models were used to study the influence of baseline neuron activity parameters (resting membrane potential and spike frequency and amplitude) on the response to SMF.

Additional analyses assessed the effect of ambient light in the laboratory on the patch-clamp recordings, and data acquired during high ambient illumination were excluded from the main analyses, as detailed in the Supplementary Material.

## Results

### SMF acutely increases sEPSC frequency

sEPSC recordings were made during 10 min SMF or sham exposure, as well as during 6 min SMF exposure (Fig. 2). There were no significant baseline differences among the neurons for these three stimulation conditions in resting membrane potential as well as baseline sEPSC frequency and amplitude (F_2,47_ ≤ 1.88, p ≥ 0.164; see Table 1). In the 10 min exposure data, there was a significant main effect of time (F_10,470_ = 2.87, p = 0.0018) as well as an interaction between stimulation condition (SMF or sham) and time (F_10,470_ = 2.97, p = 0.0012) (Fig. 2B). The presence of SMF induced a significant increase in the sEPSC frequency compared to baseline between 2 and 12 min after SMF onset (t > 3.28, p ≤ 0.0107), whereas no significant change in sEPSC frequency relative to baseline was observed during sham exposure. Moreover, there was a significant difference in sEPSC frequency between SMF and sham exposure for the time points between 2 and 12 min as well as at 16 min (t > 2.51, p ≤ 0.0226). Similarly, there was a significant effect of time on sEPSC frequency during 6 min SMF exposure (F_6,58_ = 3.35, p = 0.0067), with a significant difference from baseline at 6 min post SMF application (t = 2.85, p = 0.0343) (Fig. 2C). For both duration conditions, the frequency increase appears time-locked with the application of SMF, with the effect building up within approximately 2 min and decaying at a somewhat slower rate, over 3–4 min. Across all neurons, the average increase in sEPSC frequency compared to baseline was 9.41% (range = [−12.0%, 26.8%]) and 6.55% (range = [−0.0327%, 12.4%]) during 10 min and 6 min SMF exposure, respectively. During the 10 min SMF exposure, 26 out of 29 neurons exhibited an average sEPSC frequency increase, and that was the case for 7 out of 8 neurons for the 6 min exposure (individual data points for each cell are shown in the Supplementary Material). In contrast, sham exposure resulted in average sEPSC frequency change of −1.69% (range = [−14.4%, 8.63%]), with the number of neurons exhibiting an increase corresponding to chance level (6 out of 13). There were no significant main or interaction effects of the resting membrane potential or baseline spike amplitude or frequency on the average sEPSC frequency change during SMF exposure. Upon removal of the SMF the average sEPSC frequencies returned to levels that were not significantly different from baseline.

**Figure 2.**
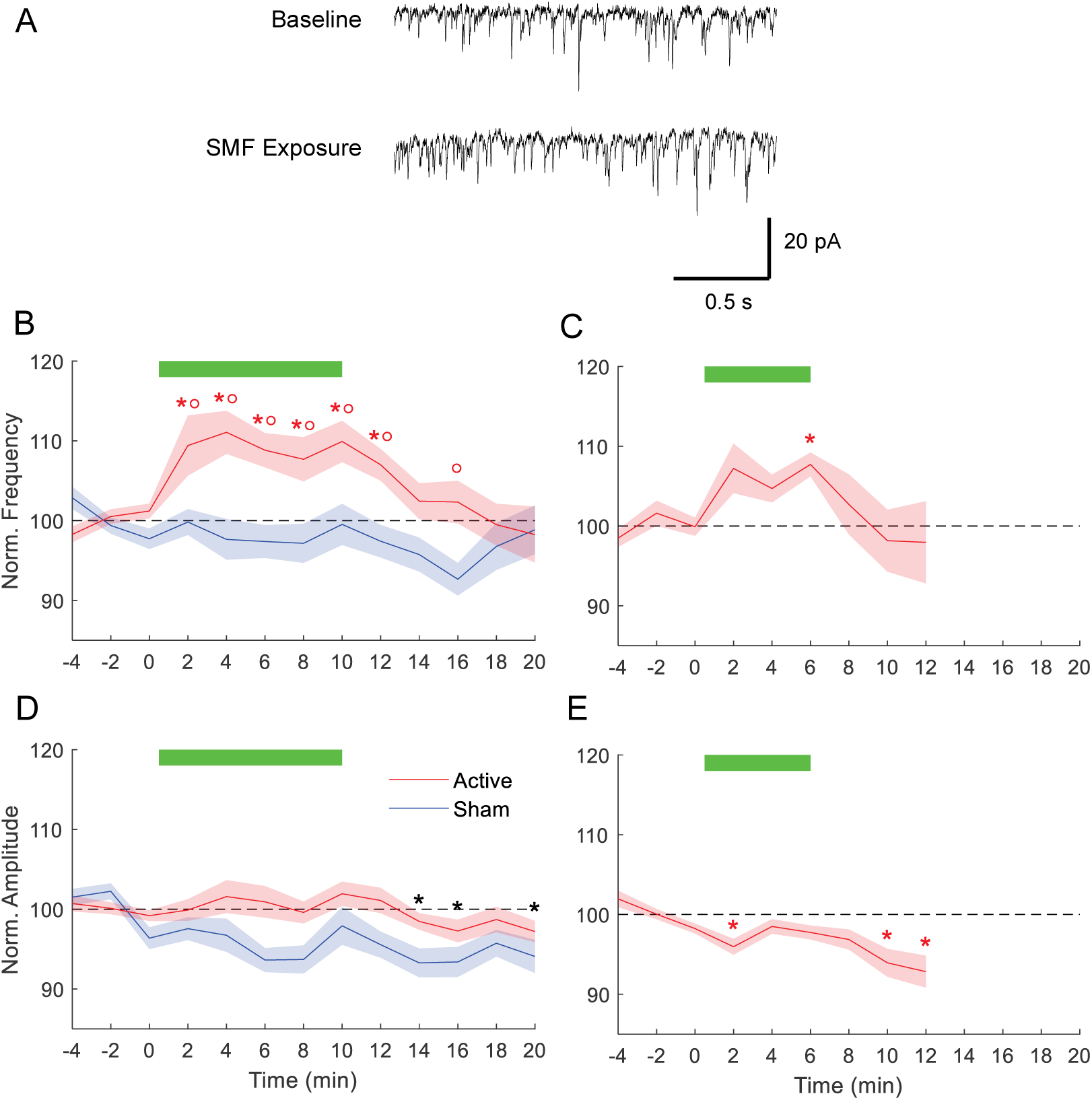
Response of spontaneous excitatory postsynaptic current (sEPSC) to SMF exposure with two durations (10 min and 6 min) as well as 10 min sham SMF. A: Example sEPSC tracings for baseline and during SMF exposure. B, D: sEPSC frequency and amplitude change relative to average baseline (normalized to 100%) during 10 min SMF (n = 29) and sham (n = 13). C, E: sEPSC frequency and amplitude change during 6 min SMF (n = 8). Thick green horizontal bars indicate presence of SMF or sham magnet. Signal time course is shown with solid line surrounded by shaded area indicating mean and standard error, respectively. The data are binned and averaged in 2 min steps. Asterisks (*) denote time points with signal significantly different from the respective baseline. Colored asterisks refer respectively to SMF or sham, whereas black asterisks refer to a main effect of time where there is no significant interaction between the two exposure conditions. Circles (°) indicate time points with significant difference between SMF and sham. Significance cutoff is p < 0.05. The individual data points are shown in Supplementary Figure S2.

**Table 1.**
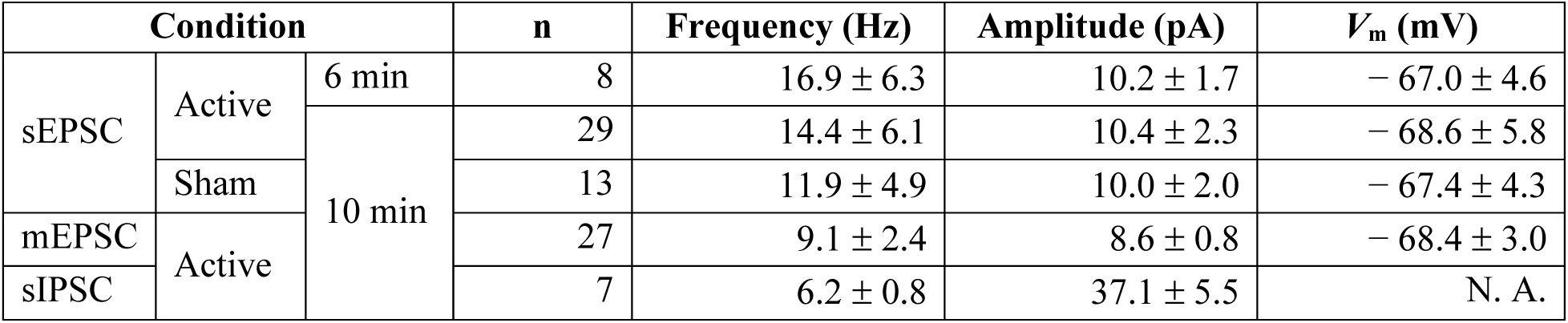
Number of cells, baseline frequency and amplitude of post-synaptic current spikes, and membrane resting potential (*V*_m_) for the experimental conditions used in the analyses. All values are mean ± standard deviation.

There was no effect of SMF on sEPSC amplitude compared to sham. For both exposure conditions there was, however, a main effect of time on sEPSC amplitude: The sEPSC amplitude gradually and consistently decreased over the course of the experiment for both the 10 min exposure condition (F_10,466_ = 3.46, p = 0.0002) and the 6 min exposure condition (F_6,58_ = 3.45, p = 0.0067). During the 10 min exposure, the sEPSC amplitude had decreased on average by 4.61% relative to baseline by 20 min after exposure onset, and during the 6 min exposure, the amplitude reduction was 7.15% by 12 min. This decrease can be explained by a slight increase in access resistance over the course of the recording, although it did not increase by more than 10%.

### SMF acutely increases mEPSC frequency

The effect of SMF exposure on mEPSCs recorded in the presence of TTX is illustrated in Fig. 3. As expected, the baseline mEPSC frequency and amplitude were significantly lower than those for sEPSCs (t ≥ 1.79, p ≤ 0.0064), whereas the baseline resting membrane potential did not differ. Analogously to the effect on sEPSCs, SMF exposure resulted in a significant increase of mEPSC frequency compared to baseline (F_10,277_ = 6.00, p < 0.0001; 2–12 min, t ≥ 3.11, p ≤ 0.0196) during the 10 min SMF exposure. The average mEPSC frequency increase was 5.39% (range = [−9.59%, 19.0%]), with 21 out of 27 neurons exhibiting an increase. There were no significant main or interaction effects of the resting membrane potential or baseline spike amplitude or frequency on the average mEPSC frequency change during SMF exposure. The mEPSC amplitude followed a declining pattern similar to that observed for sEPSCs (F_10,278_ = 9.05, p < 0.0001), reaching a mean reduction of 4.61% by 20 min after stimulus onset.

**Figure 3.**
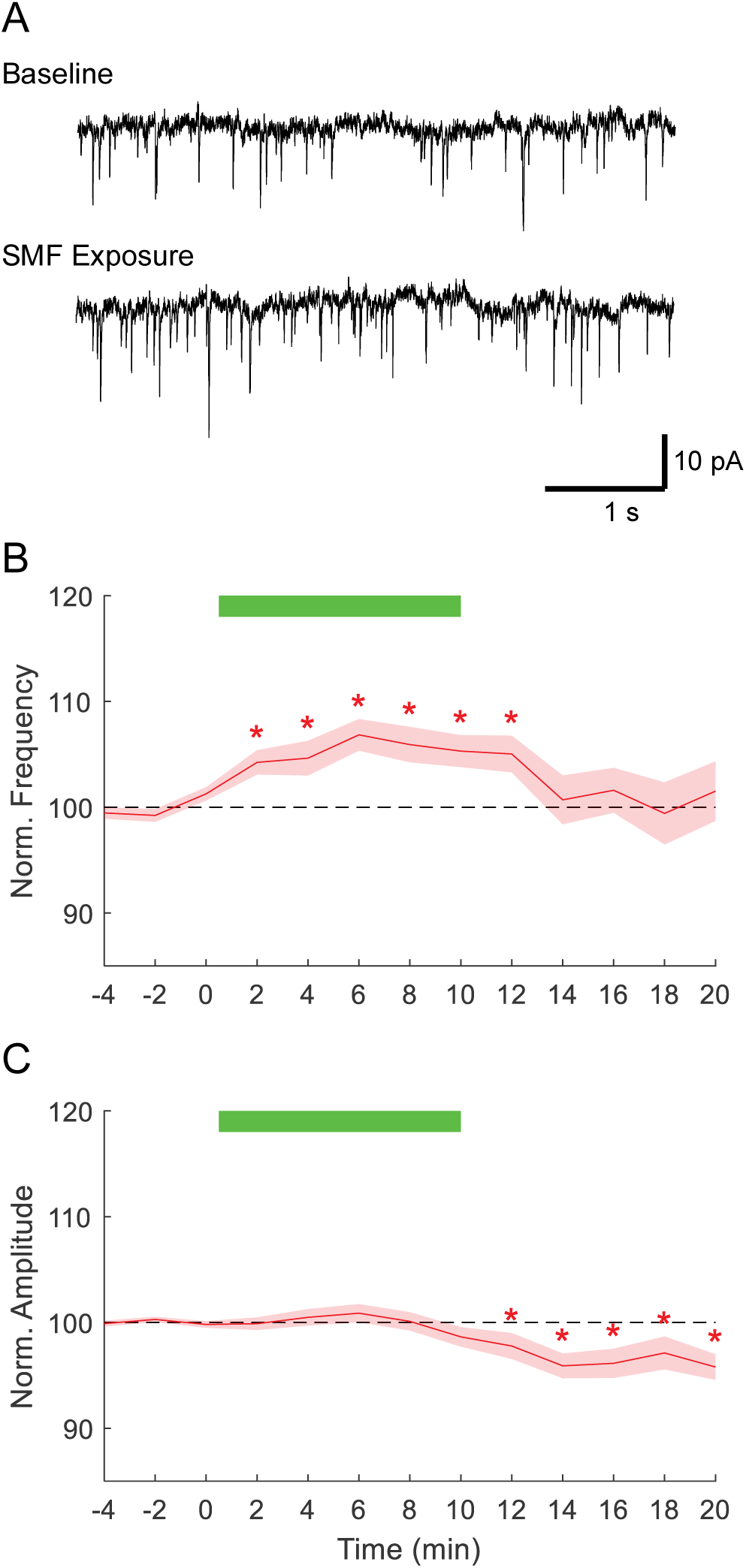
Response of miniature spontaneous excitatory postsynaptic current (mEPSC) to 10 min SMF in the presence of TTX. A: Example mEPSC tracings for baseline and during SMF exposure. B, C: mEPSC frequency and amplitude change relative to baseline (n = 27). Plot conventions are the same as in Fig. 2. The individual data points are shown in Supplementary Figure S3.

### SMF effect on sIPSCs

SMF exposure of 10 min did not affect the frequency of sIPSCs (F_10,66.2_ = 0.256, p = 0.988) (Fig. 4). Across cells the average sIPSC frequency change during the 10 min SMF exposure was only 0.853% (range = [−7.78%, 7.30%]). Like in the sEPSC and mEPSC recordings, the sIPSC amplitude decreased significantly over time (F_10,67.4_ = 2.68, p = 0.0080), declining by 10.9% below baseline at 20 min after the SMF onset.

**Figure 4.**
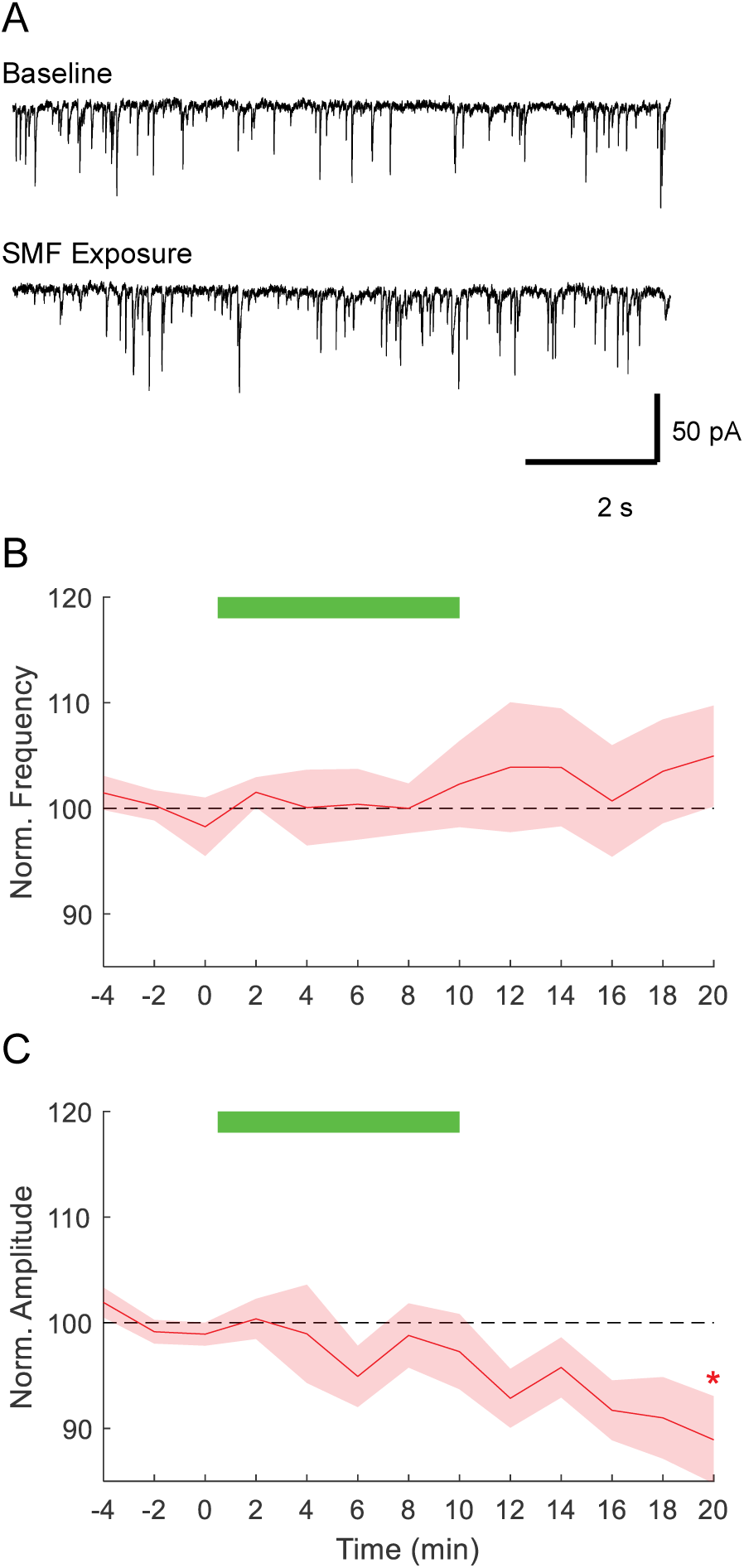
Response of spontaneous inhibitory postsynaptic current (sIPSC) to 10 min SMF. A: sIPSC tracings for baseline and during SMF. B, C: sIPSC frequency and amplitude change relative to baseline (n = 7). Plot conventions are the same as in Fig. 2. The individual data points are shown in Supplementary Figure S4.

## Discussion

We documented a significant acute positive and transient effect of SMF on the frequency of sEPSCs and mEPSCs in motor cortex pyramidal neurons in rat brain slices. SMF reached a flux density of 0.2–0.35 T in the brain slices, which overlaps with the estimated field strength of previously reported SMF stimulation of human cortex (0.12–0.25 T) (61, 62). The SMF affected sEPSC and mEPSC frequency but did not have an identifiable effect on their amplitude. This suggests a presynaptic or synaptic source of the effect if we consider that the frequency reflects the postsynaptic detection of released excitatory neurotransmitter waves, whereas a postsynaptic effect would mostly affect the ionic current, and hence amplitude, of the response to each wave. The consistency of the observed effect between sEPSCs and mEPSCs suggests that it is not due to SMF exposure influencing action potential propagation in the presynaptic axon.

A synaptic source of the SMF effect would be consistent with reports in the literature that demonstrated the presence of SMF neuromodulation in the human motor cortex only for probing with stimuli that primarily activate presynaptic stages of pyramidal motor neurons (1). Thus, SMF exposure of the human cortex appears to also affect synapses or mechanisms that involve synapses. Further, this observation would be consistent with the studies of Rosen et al. who reported SMF modulation of acetylcholine release in the mouse neuromuscular junction (63).

The significant effect of SMF on sEPSCs as well as mEPSCs but not on sIPSC and the evidence of a presynaptic effect may indicate that the signal source is predominantly located in the relatively long excitatory fibers synapsing onto the layer II/III pyramidal cells versus the relatively short, local inhibitory input fibers.

Our work found acute excitatory effects in postsynaptic recordings during 10 minutes of SMF exposure in rat brain slices but did not detect subsequent effects up to the end or recording 10 minutes post SMF. In comparison, Sinha et al. found inhibitory effects after longer, 30 minute SMF exposure with comparable field strength in pyramidal cells using whole-cell patch clamp recordings from mouse brain slices (55), in line with the studies of SMF applied transcranially to the human cortex (1, 6). They did not record the cell responses during the SMF exposure, and the subsequent inhibitory effects appear to have a different mechanism than the acute excitatory effects of our experiments. The post-exposure effect appeared to be linked to increased chloride conductance as an intrinsic modulation effect. Sinha et al. also applied TTX during SMF exposure, but washed it out for subsequent effect detection; from the disappearance of the inhibitory effect under TTX, they concluded that it may further involve sodium channels, through the electrochemical driving force of sodium ions. The authors further blocked AMPA, NMDA, and GABA receptors, which did not eliminate the SMF effect. This appears to rule out synaptic effects, which we suspect mediated the acute effects in our study. Nonetheless, the results of Sinha et al. and our measurements do not contradict each other but rather appear to represent two different effects at distinct time scales—a post-exposure intrinsic, nonsynaptic transient inhibitory effect in Sinha et al. versus a (pre-)synaptic acute excitatory effect during SMF exposure in our report, which indicates a larger calcium-dependent neurotransmitter release. The Sinha et al. effects likely do not represent our (pre-)synaptic source as it would vanish under TTX, which was not the case in our study. The increased SMF duration compared to our work can be a significant contributing factor as studies in humans have shown a more pronounced inhibitory effect for longer transcranial SMF exposure, which was also associated with different intracortical excitability changes (1, 6). However, the mechanisms underlying the effects in both Sinha et al. and our study may involve ion-channel effects, potentially through an interaction with the membrane. Indeed, Sinha et al. also reported transient swelling of the cells exposed to SMF (55). One possible membrane-mediated mechanism involves alignment of lipid rafts that, in turn, modulates the state of associated TRP channels (38).

The increased sEPSC frequency indicates an excitatory effect on the local neuron level. Since the sIPSC frequency did not change, a local excitatory effect as a result of disinhibition does not appear likely. Thus, with the inhibition unchanged, SMF appeared to cause a genuine excitatory effect. As the effect persisted under TTX, the observations cannot be attributed to upstream or more complicated network effects either. However, if an SMF effect on presynaptic calcium is involved, it may be able to drive both excitation and inhibition dependent on the affected neurons. Indeed, both intracortical inhibition and facilitation can be enhanced or suppressed depending on the parameters in transcranial SMF studies (1, 6). Furthermore, the local excitatory effect may drive a broader network inhibitory effect in the cortex.

The sEPSC frequency increase built up within 2 min or faster after onset of SMF exposure. Our observation granularity was 2 min due to the necessary averaging to manage neural variability. The time scale we observed is consistent with mouse neuromuscular junction data that showed that the effect of SMF builds up within 50–100 s (63–65). The observed dynamics implicate relatively fast phenomena such as ionic motion and membrane signaling with receptors. Typical intracellular signaling via kinases and other cytosolic enzymatic cascades subsequently sensitizing receptor or transmitter systems develops within minutes. Thus, it could potentially be involved as well. Transcriptional and epigenetic regulation, in contrast, occur over much longer time scales and are therefore not a rational explanation for the observed effect. The above-cited experiments at the mouse neuro-muscular junction reported that the effect can be abolished by removing extracellular calcium, which would indicate a calcium influx as the source of the phenomenon (63–65). Alternatively, the SMF could affect intracellular calcium ions and/or oxidative stress in astrocytes (1, 8, 51, 53, 66), leading to the release of gliotransmitters such as glutamate, which in turn modulate synaptic transmission and activity(67).

Although there was no effect of SMF on current amplitudes under any of the conditions tested, we did observe a significant main effect of time throughout the recording period. The decrease was statistically significant but gradual and small (< 10% of baseline over the course of the recording period) and did not appear to be influenced by the presence of the magnet. Potential causes for this reduction in current include changes in patch-clamp access resistance and dilution or functional loss of intracellular signaling molecules over time. It is important to note that a decrease in amplitude of less than 10% over a duration of 30 minutes is not uncommon in patch-clamp recordings of fresh brain slice preparations (68).

Collectively, our results support the interpretation that the observed effects are due to the influence of the SMF on the neural tissue and not to artifacts associated with the introduction of the magnet or other electrostatic factors associated with placing the active or sham magnet near the recording electrode. First, the magnet was moved slowly over the slice (∼ 3 °/s) to preclude the induction of significant electric current in the slice and bath. Second, the effect did not occur with sham (0 T) SMF, which rules out mechanical movement, thermal, and optical reflectance artifacts. We did detect effects of intense illumination on neural activity, and reduced the light level to mitigate this confound (see Supplementary Materials). Finally, the effect did not occur for sIPSCs and for subgroups of the neurons in which sEPSCs and mEPSCs were recorded. This observation excludes a direct effect of SMF on the patch clamp pipette. The mechanism of how the SMF is influencing excitatory synaptic signaling remain to be determined.

### Limitations and future directions

This work had several limitations. The slices were obtained from 3–5 week old male rats, a period of rapid cortical maturation which can result in neuron property variation, while not capturing the influence of sex. The placement of the slice under the edge of the permanent magnet maximized the magnetic field exposure while providing access of the pipette for patch-clamp recording, but the range of the magnetic field strength and gradient at this location (Fig. 1D) can be another source of variation. Further, the experiments were conducted at 22–25°C, well below the physiological temperature of 37°C, which affects ion diffusion, ion channel kinetics, membrane fluidity, and network dynamics, and can therefore influence related SMF mechanisms. While we did not detect a significant effect of SMF on sIPSC frequency, the sample size (n = 7) was small and had limited power to exclude moderate effects; nonetheless the average effect was small (< 1% change) and the range was within that of sham.

In order to further understand the dynamic effects of SMF within cortical layers, future studies involving the use of microelectrode arrays and cultured slices could be informative. Since the magnet is positioned above the slice, a microelectrode array design would allow the testing at various electrical stimulation as well as recording sites. Such a setup might further allow modelling of motor cortex modulation in humans with trans-synaptic excitability probing through test pulses. In addition, further studies with pharmacological manipulation of both intracellular and extracellular calcium sources are warranted to explore the contribution of this signaling cation in the SMF-induced increase in excitatory neurotransmitter release.

## Conclusions

This paper presented the first direct measurements of acute activity modulation of cortical neurons resulting from SMF exposure in the cerebral cortex. Our results are consistent with a presynaptic or synaptic site of an acute excitatory effect of SMF. These observations can inform further interrogation of SMF neuromodulation mechanisms that can ultimately aid the development or refinement of non-invasive therapeutic strategies for treating neurological and psychiatric disorders.

## Authorship Statement

**Rebecca Klein:** Conceptualization, Methodology, Investigation, Formal analysis, Visualization, Data Curation, Writing - Original Draft; **Stefan Goetz:** Conceptualization, Methodology, Investigation, Writing - Original Draft; **Wolfgang Liedtke:** Funding acquisition, Methodology, Writing - Review & Editing; **Scott Moore:** Conceptualization, Funding acquisition, Methodology, Resources, Supervision, Writing - Review & Editing; **Angel Peterchev:** Conceptualization, Funding acquisition, Methodology, Formal analysis, Visualization, Resources, Supervision, Project administration, Data Curation, Writing - Original Draft

## Data Availability

Individual data used in the table, figures, and statistical analyses are available from the Duke Research Data Repository (doi: https://doi.org/10.7924/r4r548).

## Acknowledgments

Preliminary partial results were presented at the IEEE Neural Engineering Conference, San Diego, CA, USA, 6–8 November, 2013 (69). The authors thank Drs. Warren M. Grill, Thomas J. McIntosh, David N. Beratan, Sridhar Raghavachari, Sidney A. Simon, and Leonard D. Spicer for helpful discussions.

