## Supplementary Material for "Acute Increase of Excitatory Activity in Pyramidal Neurons of Rat Motor Cortex under Static Magnetic Field"

Rebecca C. Klein[a,b], Stefan M. Goetz[a,e,f], Wolfgang B. Liedtke[c], Scott D. Moore[a,b], and Angel V. Peterchev[a,d,e,f*]

[a] Department of Psychiatry & Behavioral Sciences, Duke University, Durham, NC, USA

[b] Durham VA Medical Center, Durham, NC, USA

[c] Department of Neurology, Duke University, Durham, NC, USA

[d] Department of Biomedical Engineering, Duke University, Durham, NC, USA

[e] Department of Electrical & Computer Engineering, Duke University, Durham, NC, USA

[f] Department of Neurosurgery, Duke University, Durham, NC, USA

[*]Corresponding author:

Angel V. Peterchev

Department of Psychiatry & Behavioral Sciences

Box 3620 DUMC

Durham, NC 27710

USA

### Analysis of Impact of Ambient Light on Patch Clamp Recordings

#### Introduction

In the course of the reported study, the experimental setup was relocated from one lab (Lab 1) to another (Lab 2). In Lab 2 we noticed that sham stimulation was producing significant effects on the patch-clamp recordings of neural activity, which had not been observed in Lab 1. We noted that the overhead lights were brighter in Lab 2 compared to Lab 1. Indeed, the effect of sham stimulation disappeared when we reduced the light level in Lab 2, as described below.

In this supplement we analyze this unanticipated effect of ambient light variation on our patch clamp recordings. When the magnet or sham cylinder is moved over the slice, the light reaching the slice is reduced, creating a potential confounding effect on the static magnetic field (SMF) exposure. To address this question, we study the sham condition under different laboratory illumination levels encountered in the experiments, since sham removes any effects of the SMF. Further, we look for potential effects of the two different lab environments with low illumination on both the active and sham conditions before pooling these data together for the main analysis. We demonstrate that for the SMF exposure dataset analyzed in the main paper, the effect of light variation was insignificant. Finally, we discuss potential mechanisms of this light sensitivity.

#### Methods

Sham sEPSC data were acquired in three lab conditions: Lab 1; Lab 2, high illumination; and Lab 2, low illumination. The first two conditions used an unpainted magnet and a matched aluminum cylinder for sham. In the third condition, the magnet and sham cylinder were covered in flat black paint to minimize light reflection. Further, sham mEPSC data were collected in Lab 2, high illumination, to determine whether the effect generalized to mEPSCs. Table S1 summarizes the number of neurons recorded for each condition.

Illumination levels were measured with a photometer (Model# M110, Anaheim Scientific, Yorba Linda, CA, USA). The light intensity reaching the slice preparation in the various lab conditions and magnet configurations is reported in Table S2.

Statistical analysis followed the same methods as reported in the main paper.

**Table S1** Number of neuron recordings in each lab setup and condition.

| Condition | | | n / setup | | |
|---|---|---|---|---|---|
| | | | **Lab 1** | **Lab 2, high illum.** | **Lab 2, low illum.** |
| sEPSC | Active | 6 min | 8 | 0 | 0 |
| | | 10 min | 22 | 0 | 7 |
| | Sham | | 6 | 10 | 7 |
| mEPSC | Active | | 19 | 0 | 8 |
| | Sham | | 0 | 7 | 0 |
| sIPSC | Active | | 7 | 0 | 0 |

**Table S2** Light intensity (in units of lux) reaching the slice preparation in the various lab conditions and magnet configurations.

| **Setup** | **Magnet surface** | **Magnet position relative to Petri dish** | | |
|---|---|---|---|---|
| | | **Away** | **Partially covering** | **Fully covering** |
| Lab 1 | Nickel plated | 60 | 21* | 1.3* |
| Lab 2, high illum. | Nickel plated | 216 | 75 | 4.5 |
| Lab 2, low illum. | Flat black paint | 36 | 20 | 0 |

* Estimated from magnet away condition and relative reduction in Lab 2, high illumination condition.

#### Results

To assess the influence of the lab setup on the effect of sham exposure, we conducted mixed model analysis with time, lab setup, and their interaction as fixed effects and cell as a random effect nested within lab setup. For the sEPSC frequency, the only significant effect was the interaction between time and lab setup ($F_{20,240} = 2.64$, $p = 0.0003$). A post-hoc Dunnett test revealed that only the Lab 2 high illumination condition resulted in a significant elevation of the sEPSC frequency relative to baseline, which occurred in the last 2 minutes of sham exposure and the subsequent 4 minutes (Figs. S1 and S5). For the sEPSC amplitude, there was only a significant main effect of time ($F_{10,239} = 5.71$, $p < 0.0001$), associated with a gradual reduction over time (Figs. S1 and S5). In the Lab 2 high illumination condition, the mEPSC frequency response to sham was also significant ($F_{10,71} = 2.86$, $p = 0.0048$; Fig. S6), albeit shorter than during SMF (Fig. 3 and S3). The sham mEPSC amplitude decayed with time ($F_{10,71} = 5.56$, $p < 0.0001$; Fig. S6) as in the other conditions. The baseline neural activity was affected by the lab environment only for mEPSCs ($F_{2,582} = 16.2$, $p < 0.0001$), with the effect decreasing when illumination was lowered in Lab 2 (Fig. S7). Based on these findings, we concluded that the Lab 2 high illumination environment resulted in significant effects of light on neural activity, and therefore excluded any data from this lab setup from the analysis in the main paper.

Further, to assess whether there were differences between the remaining recordings from Lab 1 and Lab 2 low illumination, the mixed model analyses of sEPSCs (10 min exposure, SMF and sham) and mEPSCs (10 min SMF) in the main paper were first conducted including lab setup as a fixed effect. There were no significant main effects or significant interactions involving lab setup. Therefore, we pooled the data from these two lab setups in the analysis in the main paper.

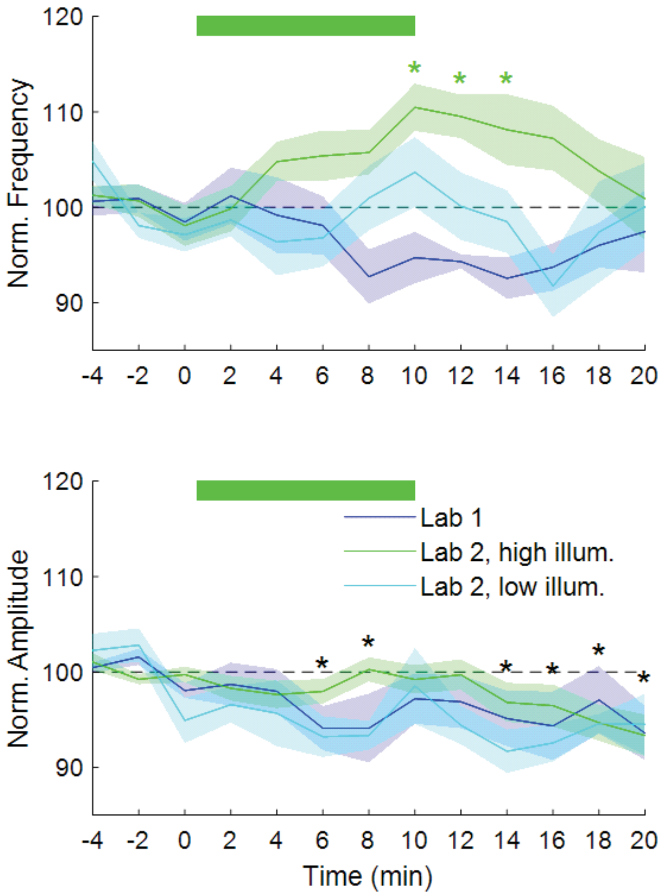


**Figure S1**
sEPSC frequency and amplitude changes associated with sham magnet exposure in three different lab environments: Lab 1 (n = 6); Lab 2 high illumination (n = 10); Lab 2, low illumination (n = 7). Signal time course is shown with solid line surrounded by shaded area indicating mean and standard error, respectively. The data are binned and averaged in 2 min steps. Relative to baseline there is a significant ($p < 0.05$) effect on frequency only in Lab 2, high illumination (green asterisks). There is a significant effect of time on amplitude (black asterisks) with no difference between conditions. Individual data points are shown in Figure S5.

#### Discussion

The above analysis suggests that the presence of relatively high intensity ambient light, as in the Lab 2 high illumination setup, can produce modulation of the recorded neuronal activity when the patch-clamped slice is covered (shaded) and then uncovered by the magnet or sham cylinder. Although the source of this effect is unclear, several known mechanisms could be contributing. First, light can affect cell metabolism (Rojas et al., 2008; Zein et al., 2018; Hayworth et al., 2010; Stockley et al., 2017; Kozai and Vazquez, 2015). Light absorption by cytochrome C oxidase appears to be a mechanism, which ultimately can lead to increased mitochondrial activity and levels of adenosine triphosphate, cyclic adenosine monophosphate, and reactive oxygen species (Yang et al., 2020; Freitas and Hamblin, 2016; Poyton and Ball, 2011; Hennessy and Hamblin, 2017; Salehpour et al., 2018). Second, measurement hardware can cause optically induced artifacts (Kozai and Vazquez, 2025). For instance, it has been reported that the Becquerel effect in patch clamp electrodes can make recorded potentials sensitive to light (Han et al., 2009). Light transients, which can correlate with a real or sham magnet moved onto a sample, may cause transient charging effects. Finally, photochemical and thermal effects can indirectly modulate neurotransmitter release (Wu and Yang, 1992; Bernard et al., 1997).

#### Conclusion

We observed an unanticipated effect of ambient light on the patch-clamp recordings of neural activity. This effect was dependent on the light intensity, and was insignificant below a certain light level. The literature suggests possible mechanisms for this phenomenon. Determining the specific mechanism of light sensitivity in our experiments requires further investigation that is beyond the scope of this paper. We conducted analysis of the data under three laboratory conditions, and in the main paper analysis we excluded data from the setup which had highest ambient illumination with significant effect on the patch-clamp recordings. Therefore, we believe that the only significant effect captured by the data in the main paper is that of the static magnetic field. These observations underscore the importance of minimizing the exposure to intense light and the inclusion of appropriate sham conditions to account for any potential effects of light.

#### Figures Showing Individual Data Points

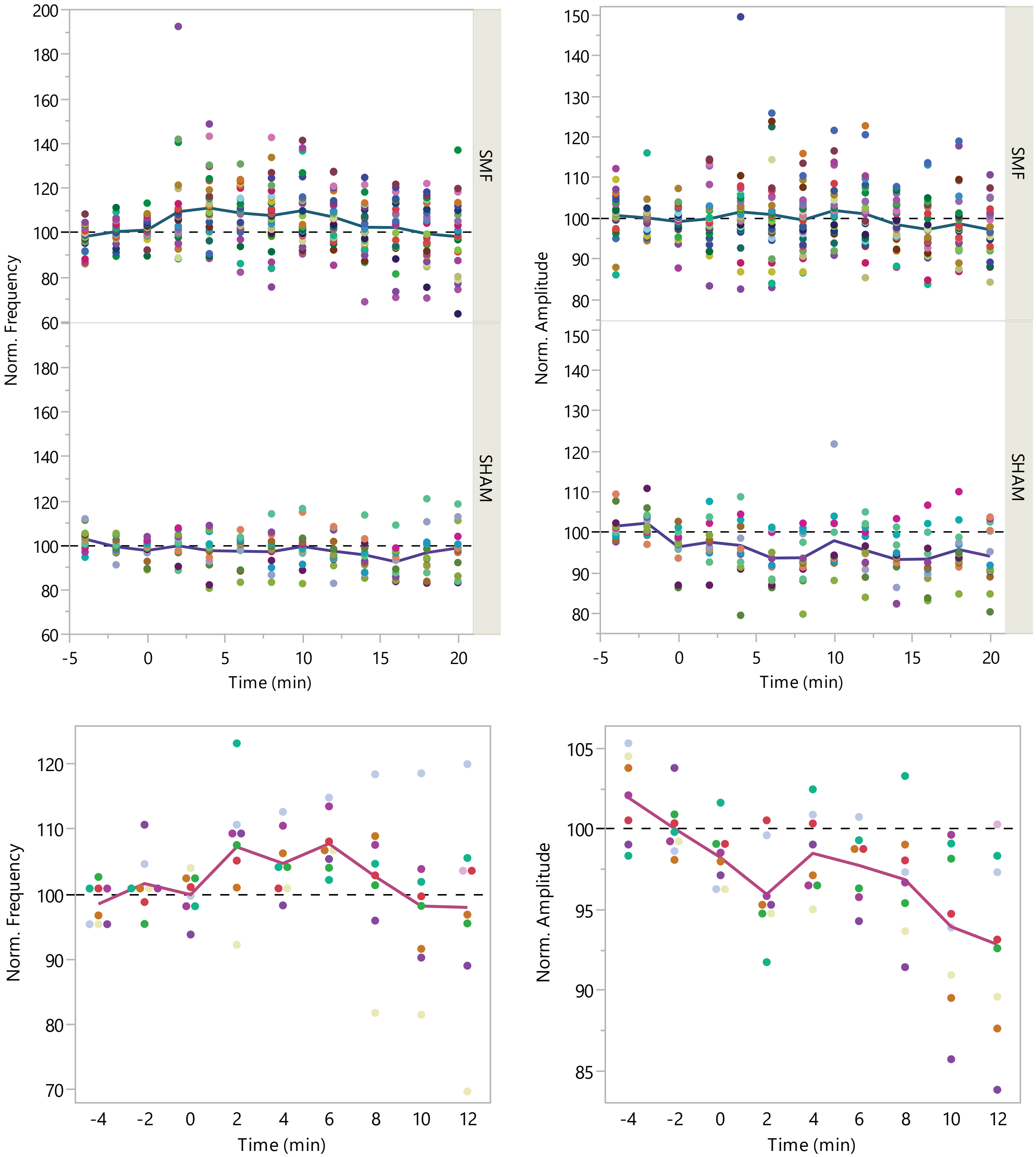


**Figure S2.** Individual data and mean for 10 min SMF and sham sEPSC in Fig. 2B,D (top) and 6 min SMF sEPSC in Fig. 2C,E (bottom).

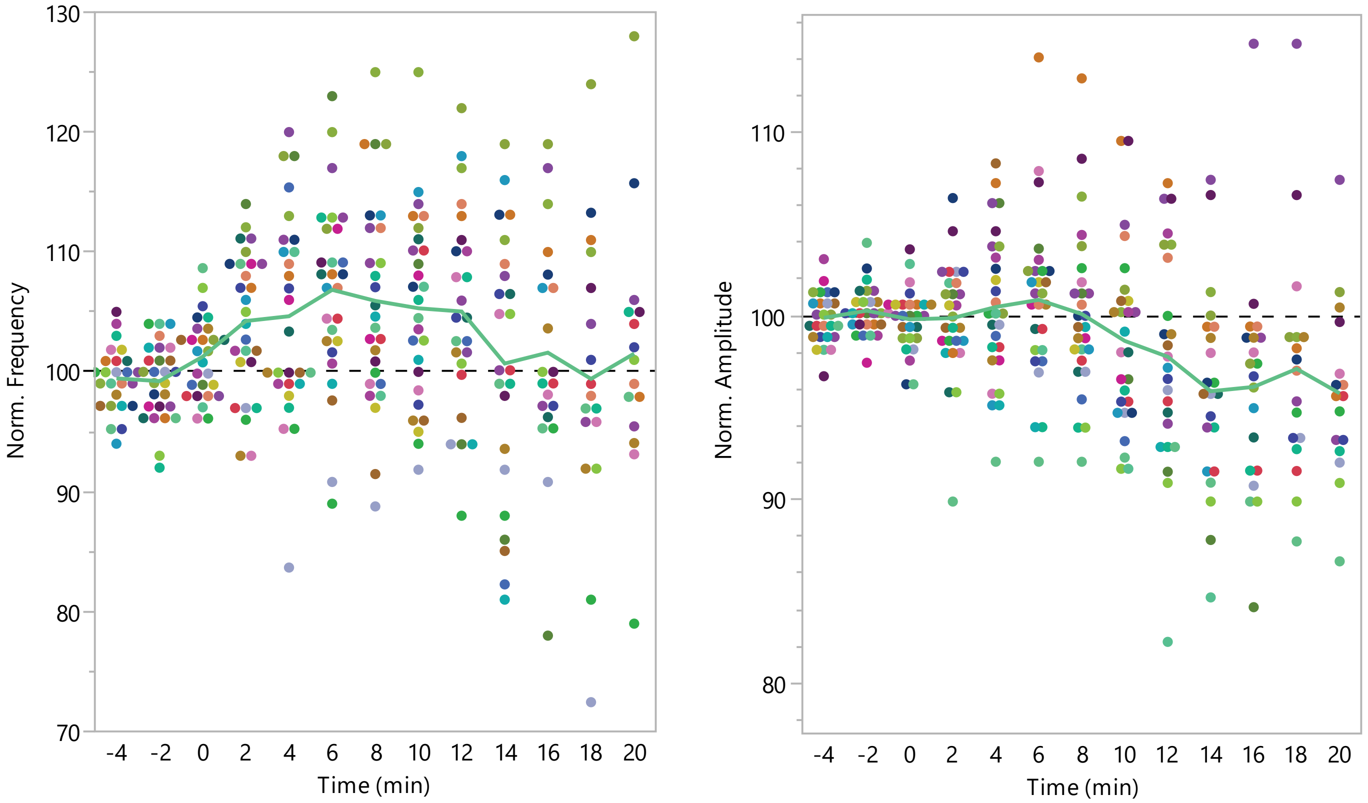


**Figure S3.** Individual data and mean for 10 min SMF mEPSC in Fig. 3B–C.

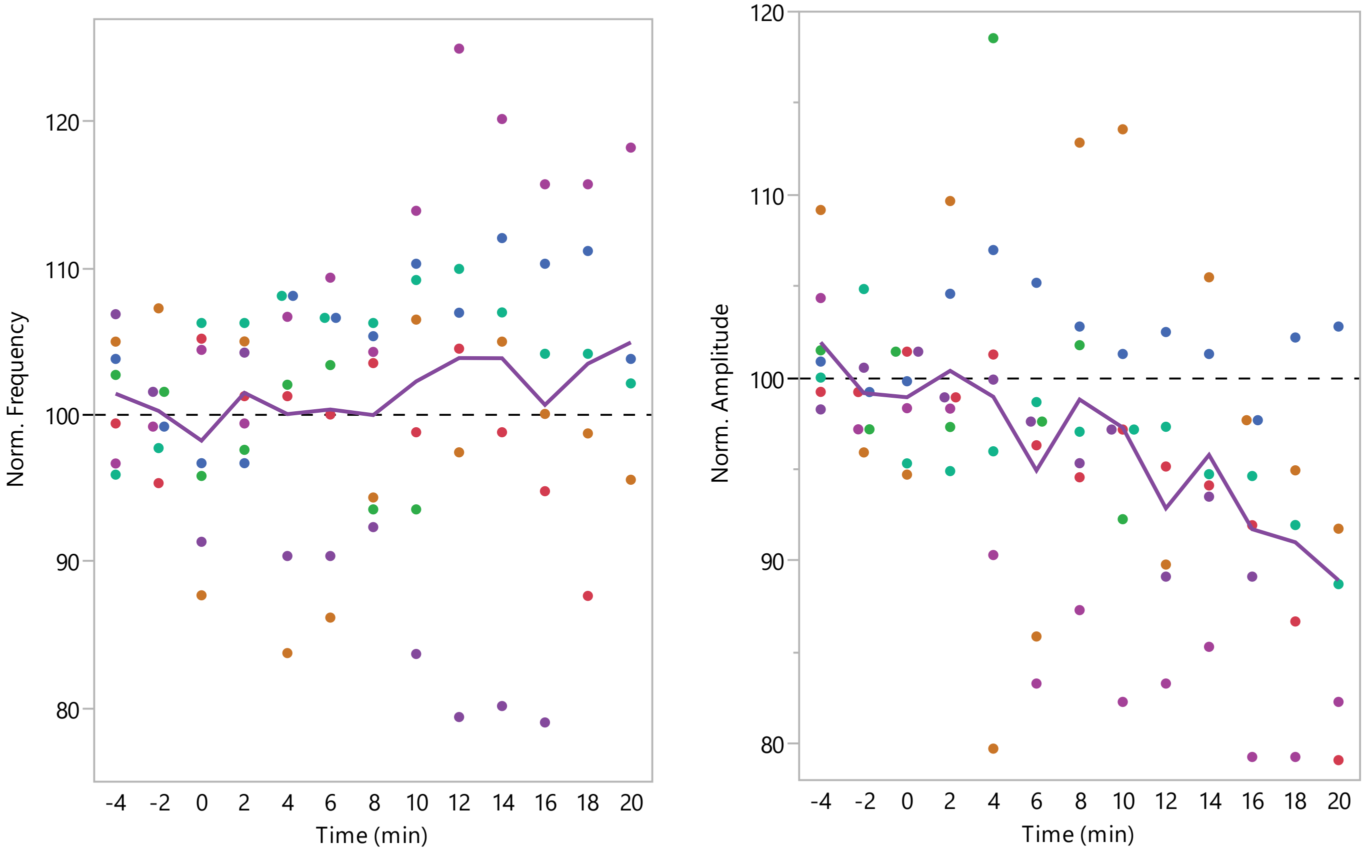


**Figure S4.** Individual data and mean for 10 min SMF sIPSC in Fig. 4B–C.

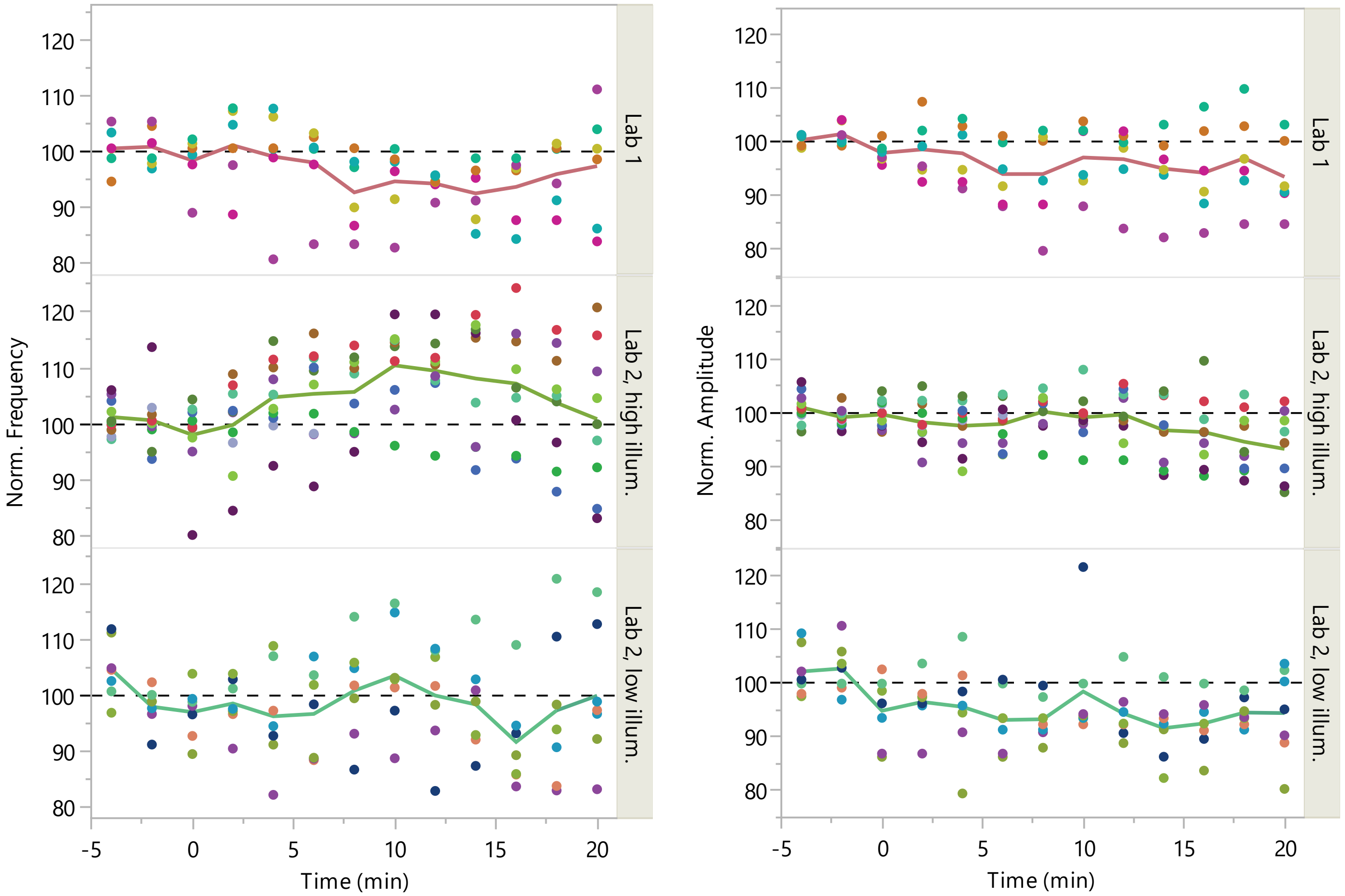


**Figure S5.** Individual data and mean for 10 min sham sEPSC in different lab environments in Fig. S1.

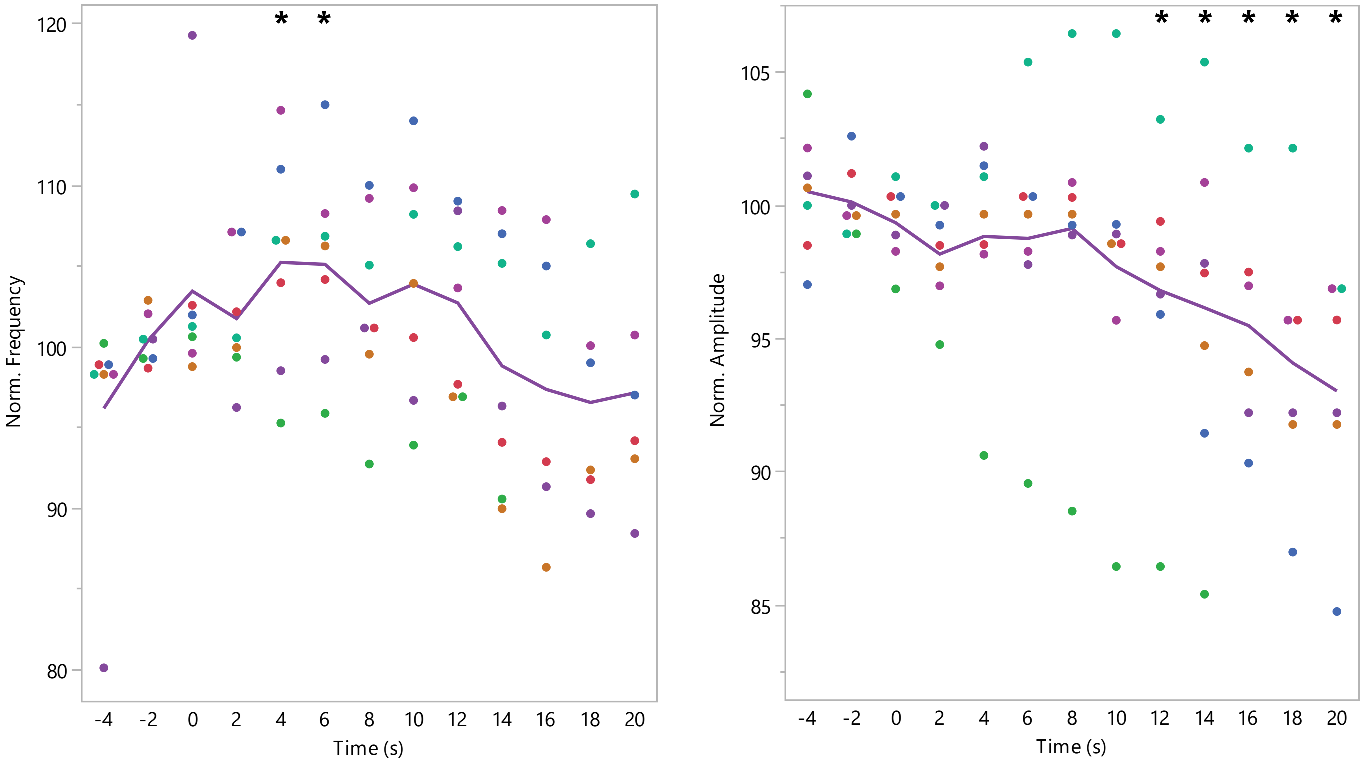


**Figure S6.** Individual data and mean for 10 min sham mEPSC in Lab 2, high illumination. * $p < 0.05$ relative to baseline (Dunnett-Hsu test).

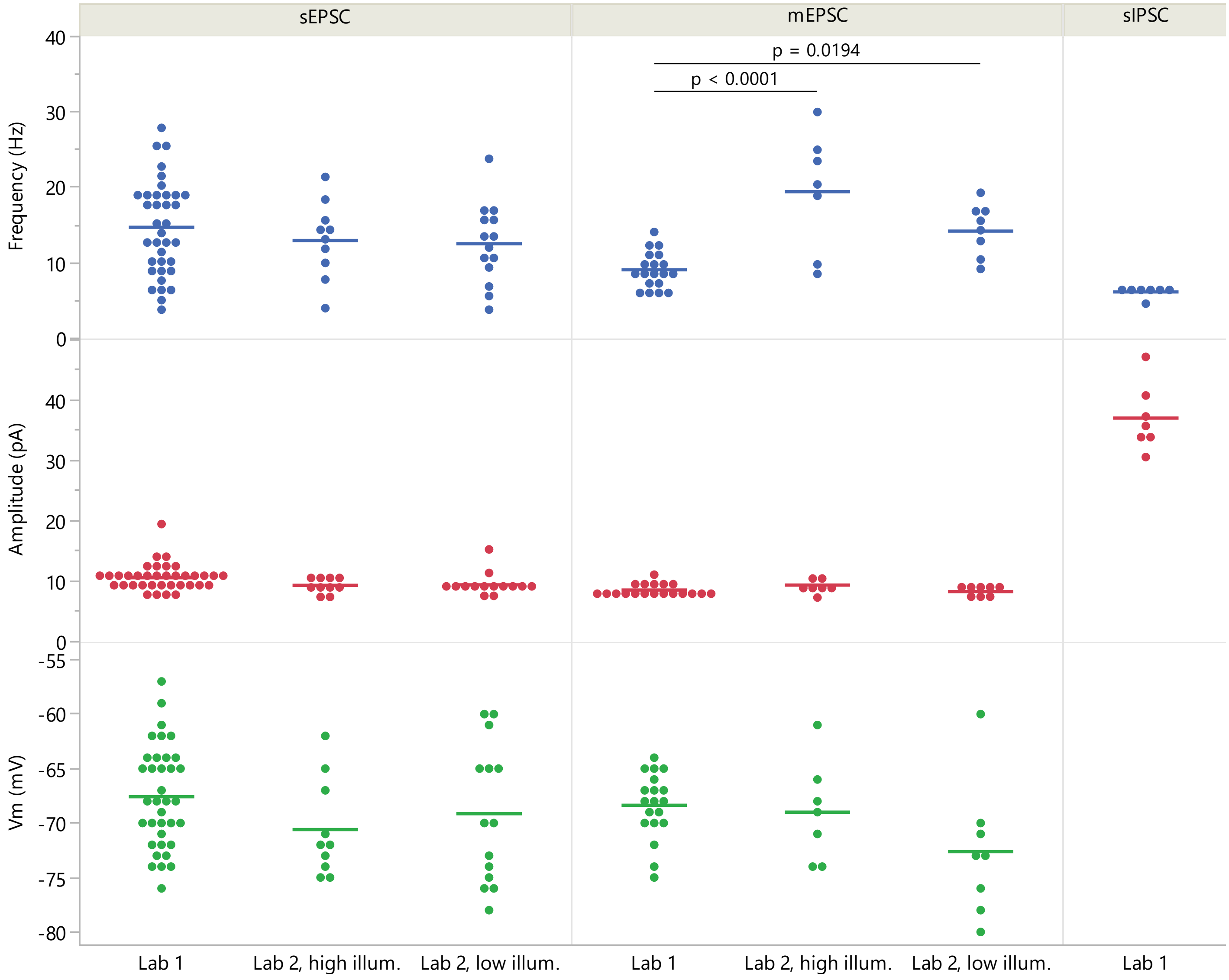


**Figure S7.** Data points for individual cells and their mean (horizontal bars) of baseline spike frequency and amplitude as well as resting membrane potential ($V_m$) for the different recording types and laboratory settings. Significant differences between laboratory settings within recording modality are indicated by their p-values.
